## Supplemental Figures 1-10 for "Structural basis of the mycobacterial stress-response RNA polymerase auto-inhibition via oligomerization"

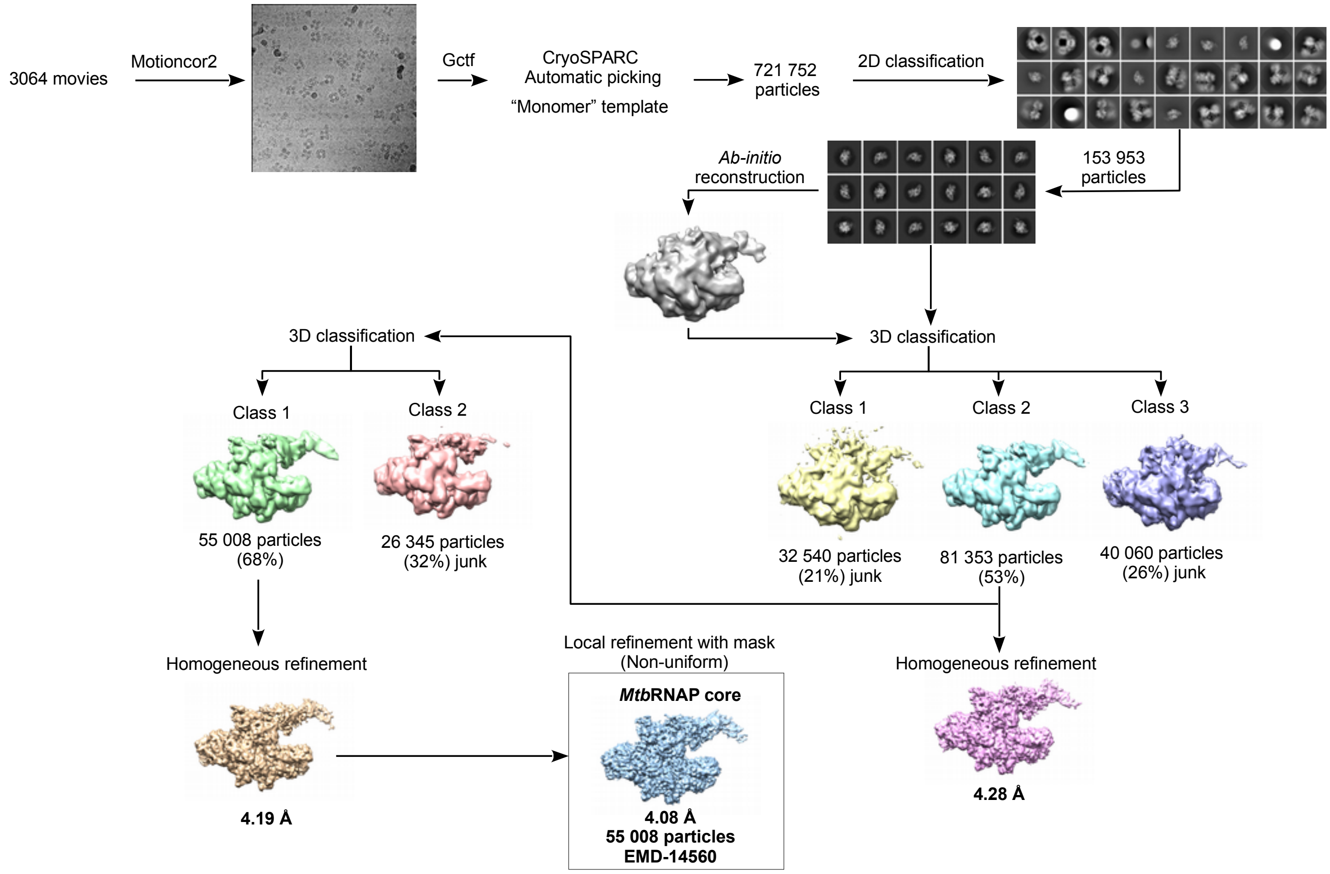


**Figure S1. The cryo-EM data processing pipeline forthe *M. tuberculosis* RNAP core**

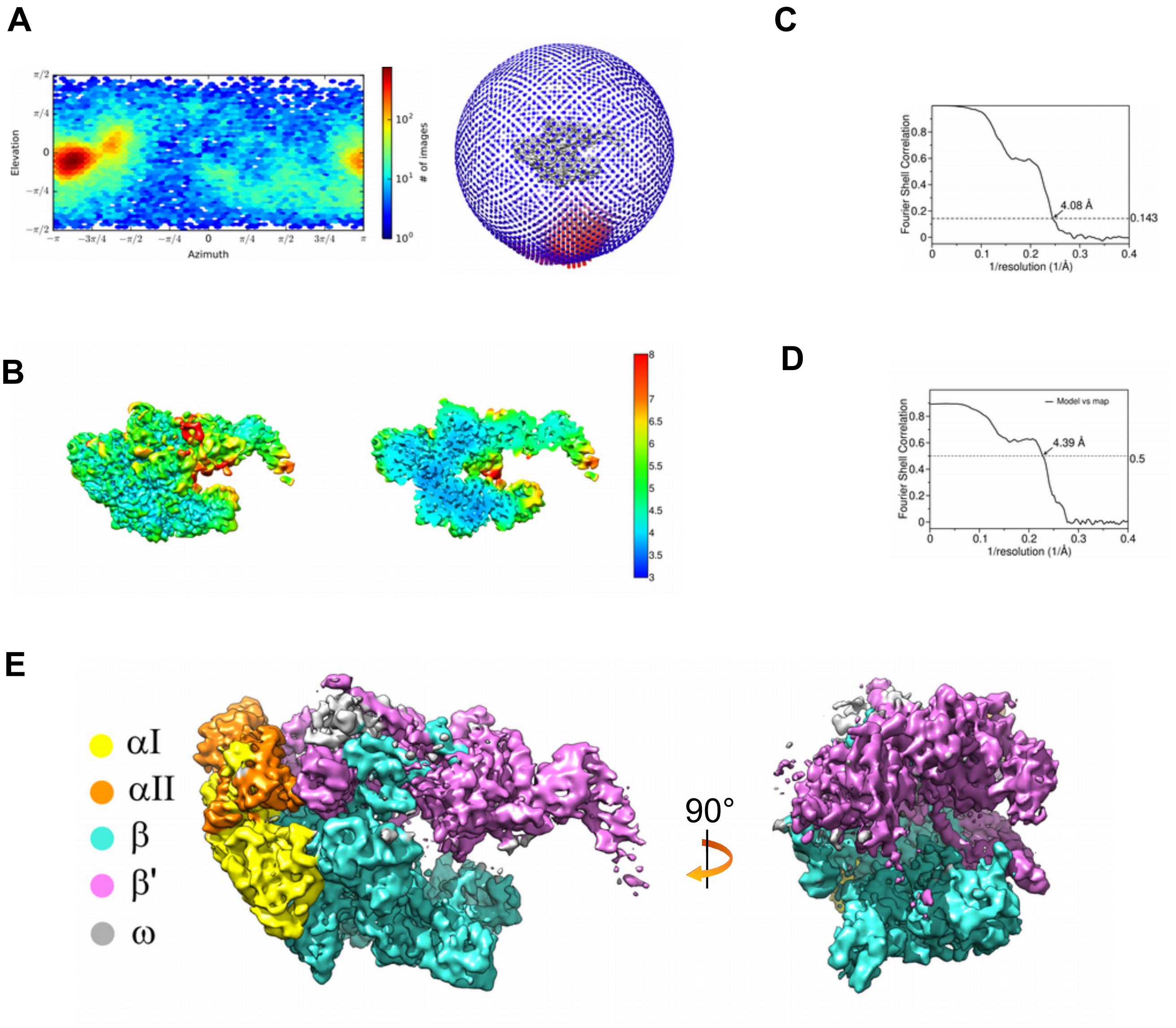


**Figure S2. The cryo-EM structure of the *M. tuberculosis* RNAP core**

(A) Heat map and spherical presentations of the angular distributions for particles projections calculated in cryoSPARC. (B) Cryo-EM density map and sliced map (on the right) colored according to the local resolution. (C) Gold-standard FSC calculated for the map in cryoSPARC. The dotted line shows the 0.143 FSC cutoff. (D) Gold-standard FSC calculated for the model vs. full-map using the MTRIAGE module of Phenix (right). The dotted line shows the 0.5 FSC cutoff. (E) Views on the cryo-EM density map of the Mtb RNAP core. The RNAP subunits are colored as indicated on the left.

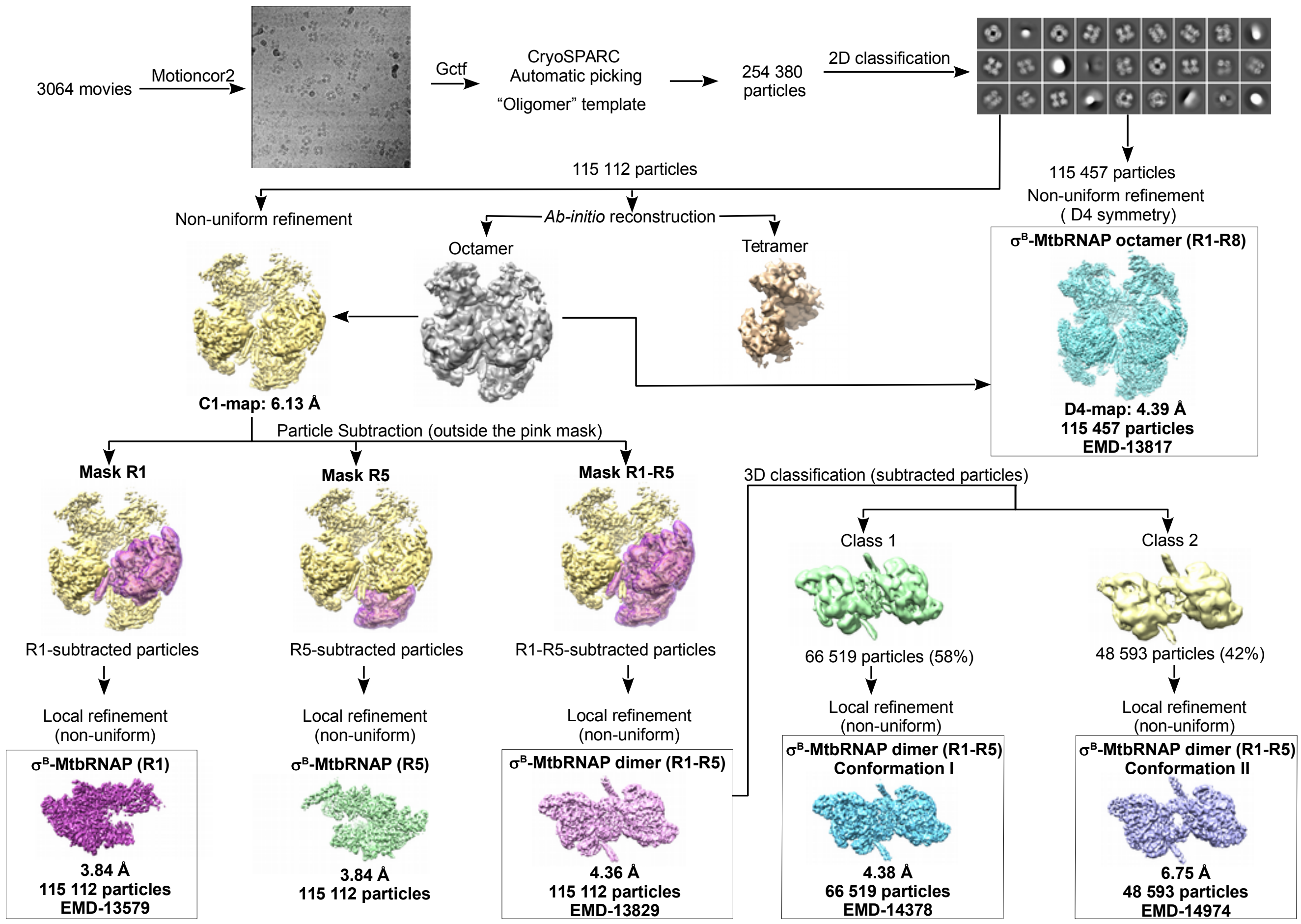


**Figure S3. The cryo-EM data processing pipeline for *M. tuberculosis* E$\sigma^B$**

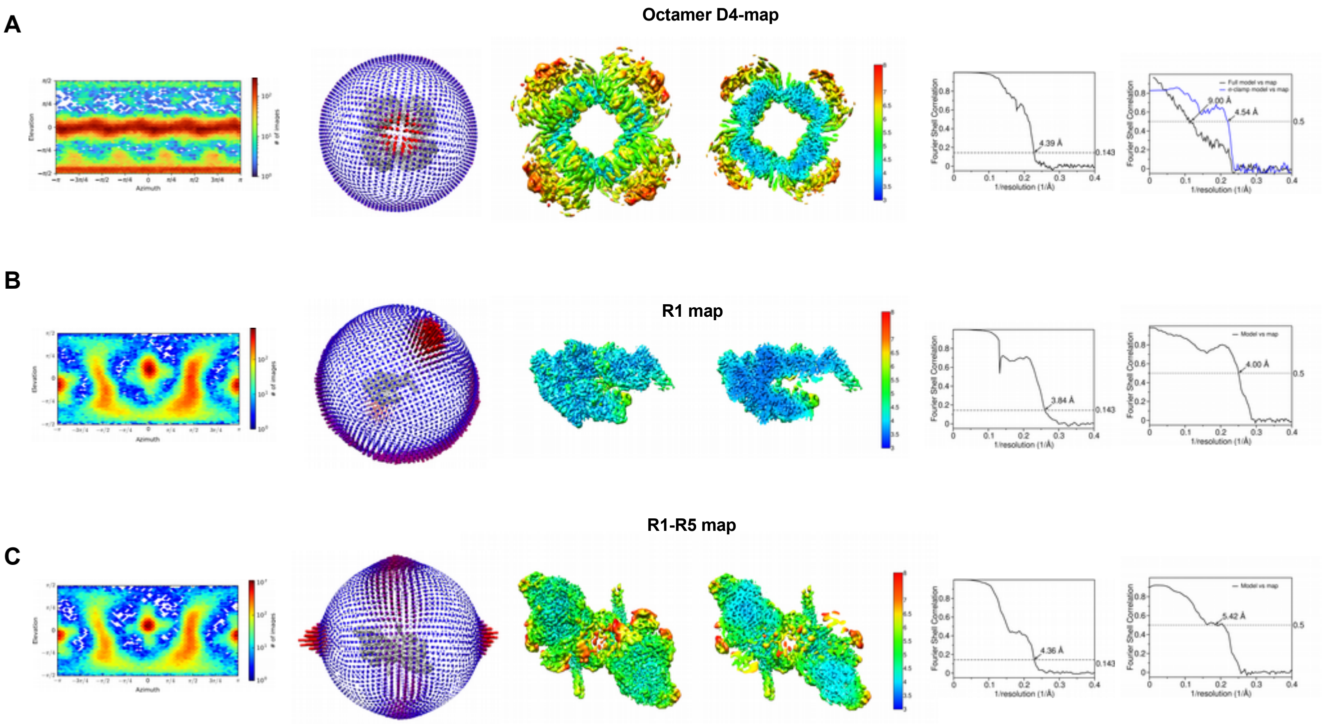


**Figure S4.** ***M. tuberculosis*** **$E\sigma^B$ octamer, monomer and dimer**

(A) $E\sigma^B$ octamer. From left to right: angular distributions for particles projections calculated in cryoSPARC and presented as a heat map and as a sphere: Cryo-EM density map and sliced map (on the right) colored according to the local resolution. Gold-standard FSC calculated for the map in cryoSPARC. The dotted line shows the 0.143 FSC cutoff. Gold-standard FSC calculated for the model vs. full-map using the MTRIAGE module of Phenix (right). The dotted line shows the 0.5 FSC cutoff. (B) $E\sigma^B$ R1 protomer. From left to right: angular distributions for particles projections calculated in cryoSPARC and presented as a heat map and as a sphere. Cryo-EM density map and sliced map (on the right) colored according to the local resolution. Gold-standard FSC calculated for the map in cryoSPARC. The dotted line shows the 0.143 FSC cutoff. Gold-standard FSC calculated for the model vs. full-map using the MTRIAGE module of Phenix (right). The dotted line shows the 0.5 FSC cutoff. (C) $E\sigma^B$ R1-R5 dimer. From left to right: angular distributions for particles projections calculated in cryoSPARC and presented as a heat map and as a sphere. Cryo-EM density map and sliced map (on the right) colored according to the local resolution. Gold-standard FSC calculated for the map in cryoSPARC. The dotted line shows the 0.143 FSC cutoff. Gold-standard FSC calculated for the model vs. full-map using the MTRIAGE module of Phenix (right). The dotted line shows the 0.5 FSC cutoff.

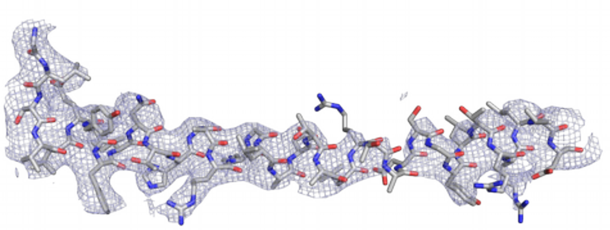

**β' bridge helix (843-879)**

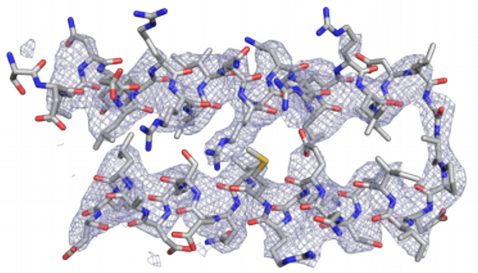

**β' coiled-coil (338-383)**

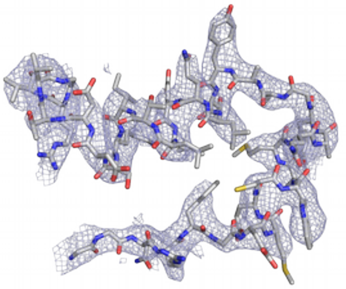

**βa15 (1064-1100)**

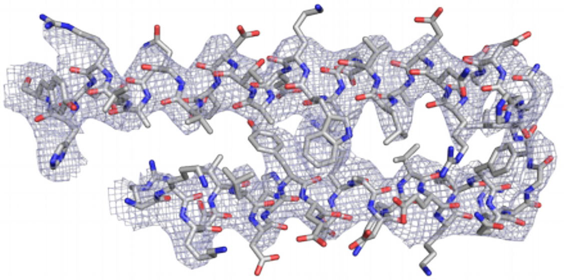

**β'a13-a14/rim helices (741-793)**

**Figure S5.** ***M. tuberculosis*** **E$\sigma^B$ R1 protomer.**

Sample density maps and molecular models of the *M. tuberculosis* E$\sigma^B$ structural elements.

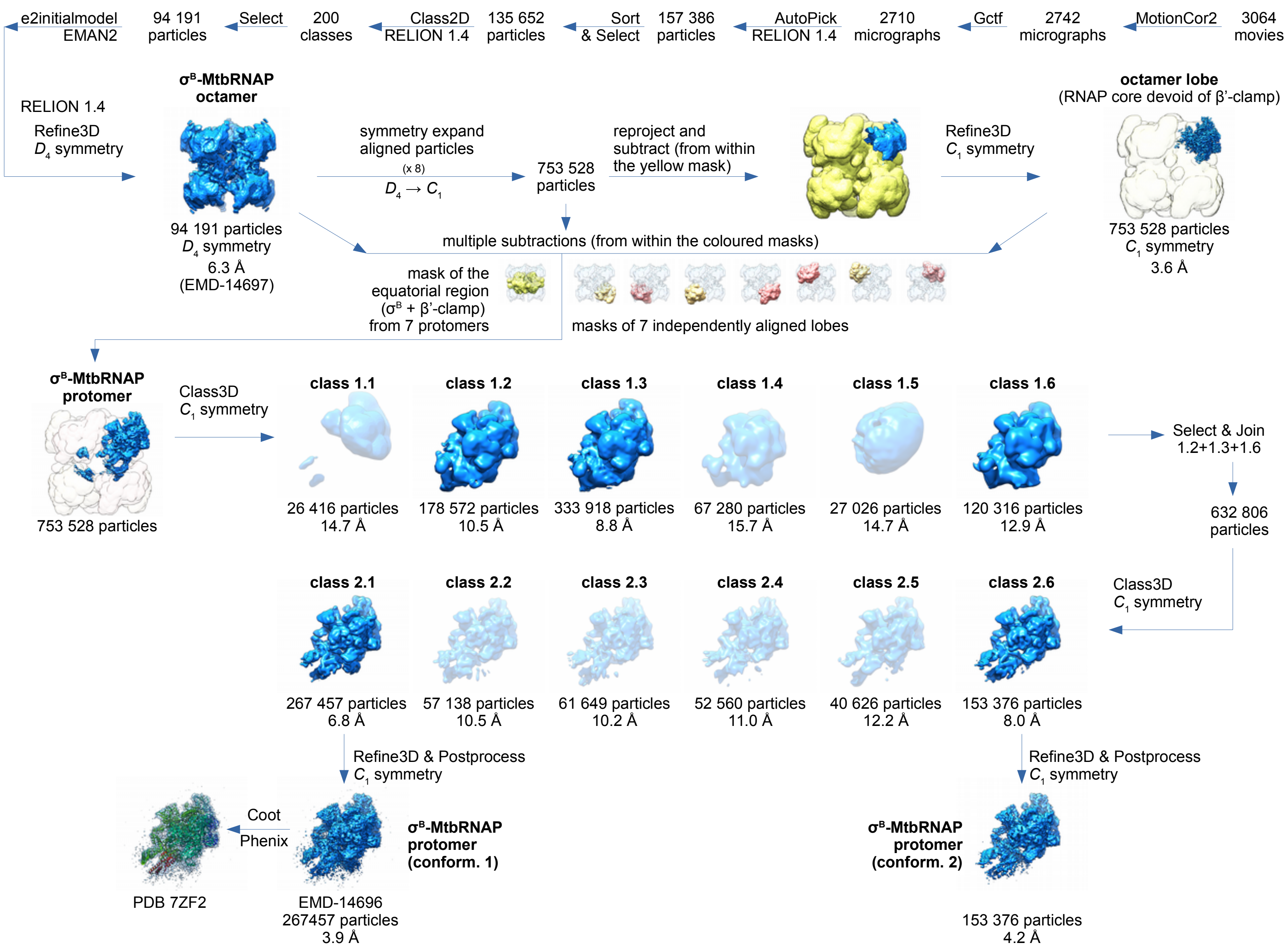


**Figure S6. Alternative cryo-EM data processing pipeline for *M. tuberculosis* Eσ$^B$ using RELION**

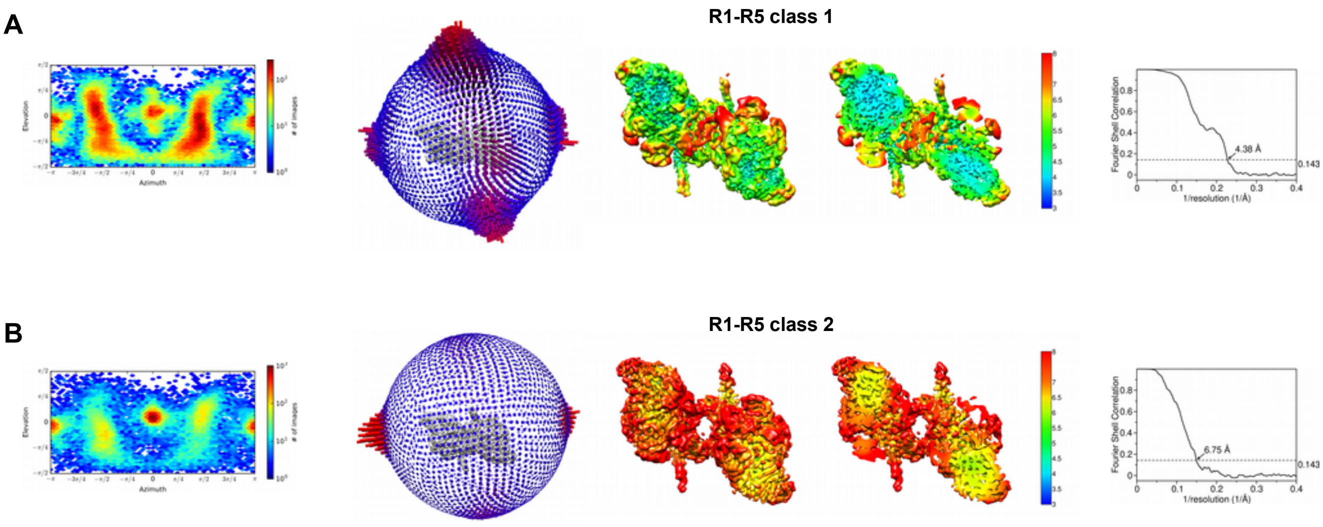


**Figure S7. Two conformations of the E$\sigma^B$ dimer**

(A) E$\sigma^B$ R1-R5 dimer conformation I (3D class 1). From left to right: angular distributions for particles projections calculated in cryoSPARC and presented as a heat map and as a sphere. Cryo-EM density map and sliced map (on the right) colored according to the local resolution. Gold-standard FSC calculated for the map in cryoSPARC. Dotted line shows the 0.143 FSC cutoff. Gold-standard FSC calculated for the model vs. full-map using the MTRIAGE module of Phenix (right). The dotted line shows the 0.5 FSC cutoff. (B) E$\sigma^B$ R1-R5 dimer conformation II (3D class 2). From left to right: angular distributions for particles projections calculated in cryoSPARC and presented as a heat map and as a sphere. Cryo-EM density map and sliced map (on the right) colored according to the local resolution. Gold-standard FSC calculated for the map in cryoSPARC. The dotted line shows the 0.143 FSC cutoff. Gold-standard FSC calculated for the model vs. full-map using the MTRIAGE module of Phenix (right). The dotted line shows the 0.5 FSC cutoff.

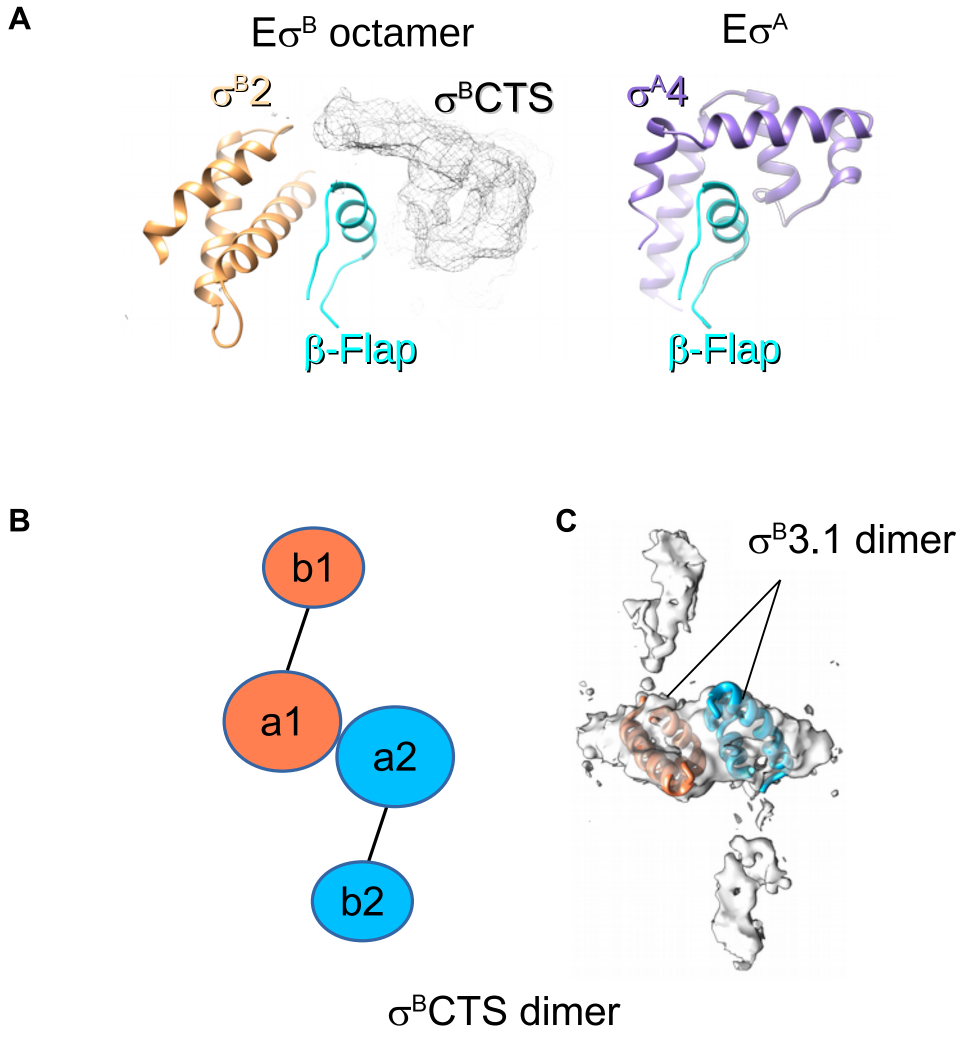


**Figure S8. The C-terminal segment of σ$^{B}$**

(A) Environment of the β flap in the Eσ$^{B}$ octamer (left) and in Eσ$^{A}$ (right). The unresolved density of the σ$^{B}$ C-terminal segment (σ$^{B}$CTS) is shown as a mesh. The σ$^{B}$ domain σ2 and the σ$^{A}$ domain σ4 are shown as ribbons. (B) Schematic representation of the two stacked σ$^{B}$ CTS in the R1-R5 (Eσ$^{B}$)$_2$ dimer. Each σ$^{B}$CTS is depicted as two connected ellipsoids. The σ$^{B}$CTS assigned to the R1 RNAP protomer is colored in coral (a1-b1) and the σ$^{B}$CTS assigned to the R5 RNAP protomer (a2-b2) is in deep sky blue. (C) Cryo-EM density of the two stacked σ$^{B}$CTS in the R1-R5 (Eσ$^{B}$)$_2$ dimer with the fitted model of the σ$^{B}$ subregion 3.1 dimer (residues 162-212, shown as ribbons). The σ$^{B}$ monomers are colored as in panel B.

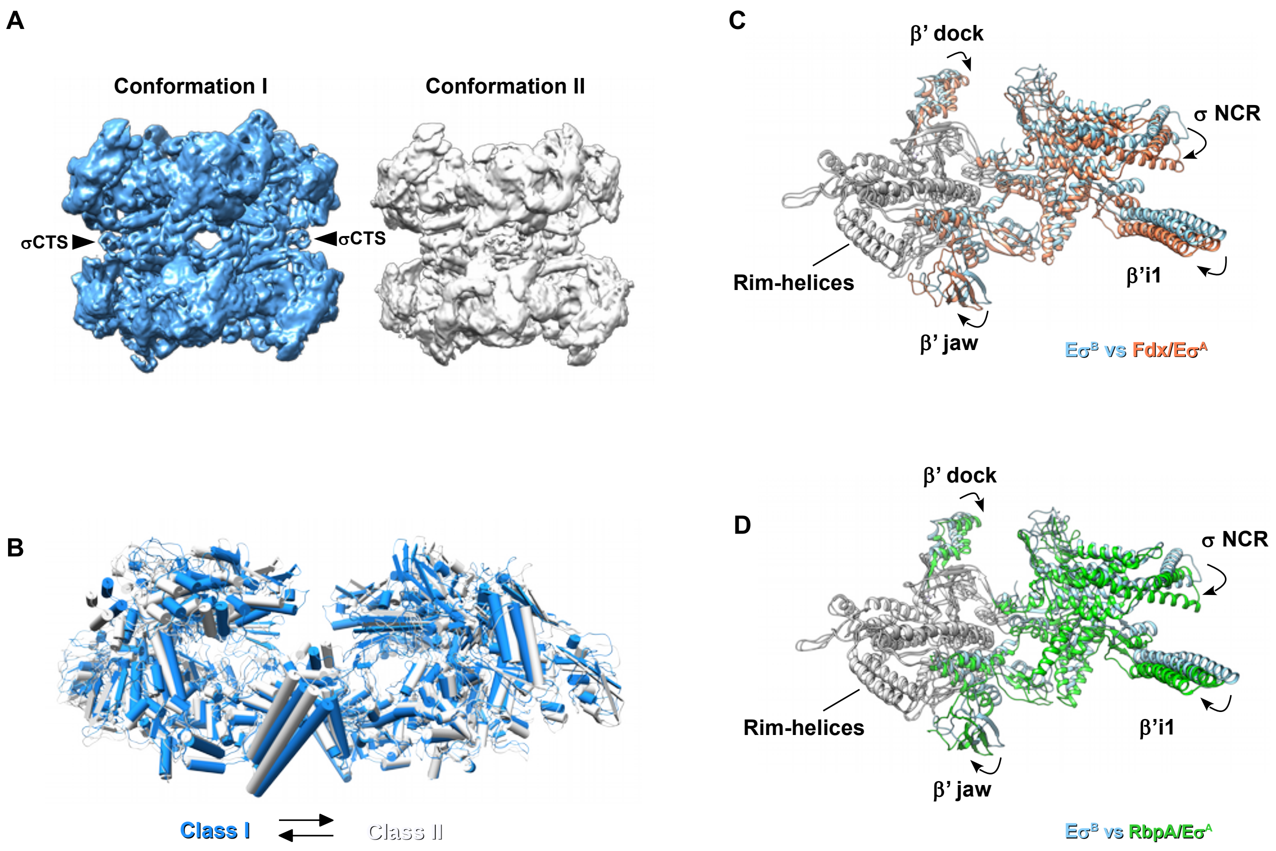


**Figure S9. Conformational changes in E$\sigma^B$.**

(A) Two conformations of the E$\sigma^B$ octamer retrieved by 3D variability analysis in cryoSPARC. The $\sigma^B$CTS positions are indicated with arrows. (B) Superposition of class I (blue) and class II RNAP (white) dimers. (C) Superposition of the β' subunit domains: β'i1 (a.a. 141-230), β'jaw (a.a. 1037-1116), β' dock (β'a11, aa 440-495) (Lane and Darst, 2010), and $\sigma^B$NCR in E$\sigma^B$ with those of E$\sigma^A$ in complex with the antibiotic Fdx (PDB ID 6FBV). (D) Superposition of the β' subunit domains: β'i1 (a.a. 141-230), β'jaw (a.a. 1037-1116), and $\sigma^B$NCR in E$\sigma^B$ with those in RbpA/E$\sigma^A$ (PDB ID 6C05).

A

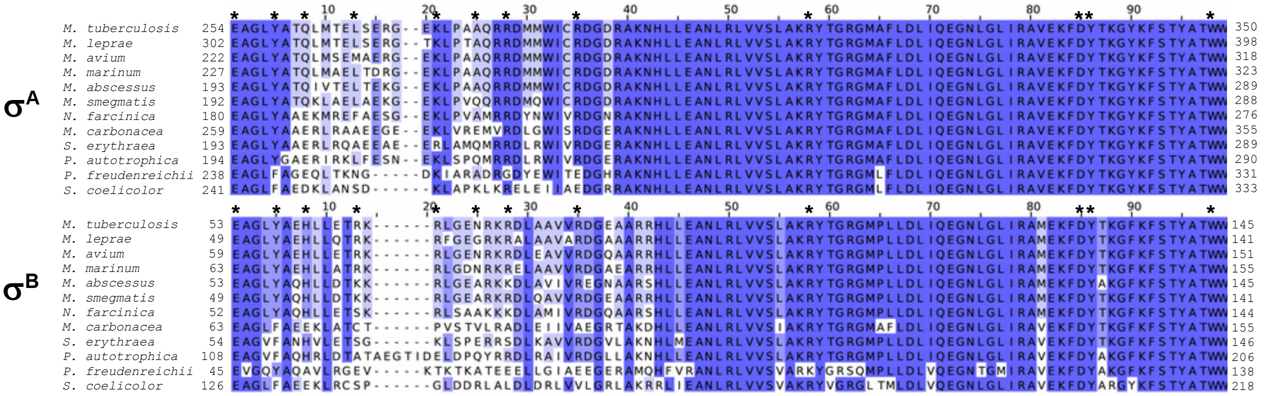


B

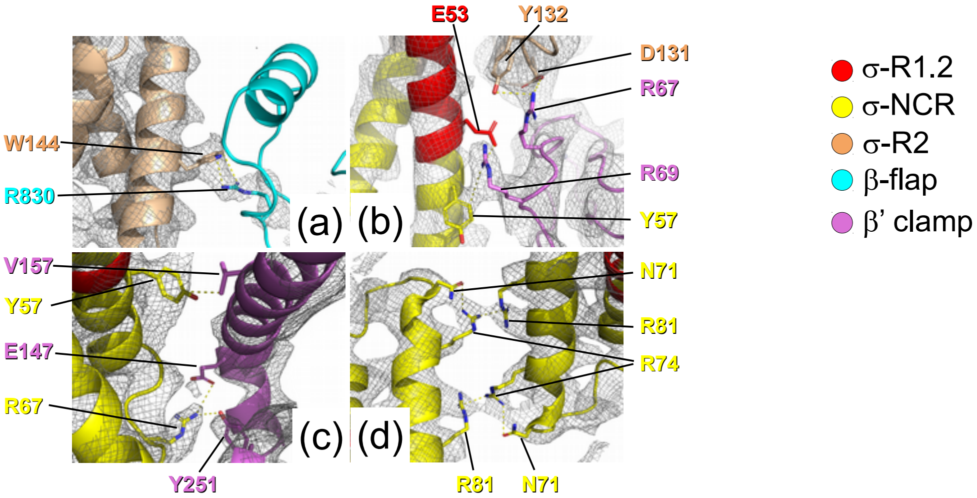


**Figure S10. Structure of the octamer-forming interfaces**

(A) Alignment of the representative sequences of the σ$^B$ and σ$^A$ subunits of *Actinobacteria* implicated in inter-face formation. Asterisks show residues implicated in inter-subunit interactions. Residues are shaded blue ac-cording to the blosum62 information score. (B) Views on the inter-subunit interfaces: (a) β flap - σ$^B$, (b) β'ZBD - σ$^B$ (c) β'i1 - σ$^B$, (d) σ$^B$NCR - σ$^B$NCR. The cryo-EM density is shown as a gray mesh with the superimposed rib-bons color coded as indicated on the right. Interacting residues are shown as sticks and labeled.
